# Dual language experience enhances neural activation variability during an fMRI reading and language task

**DOI:** 10.1101/502625

**Authors:** J.G. Malins, H. D’Silva, G. Luk, A.E. Hernandez, S.J. Frost, K.R. Pugh, W.E. Mencl, C. Mehta, J. Bosson-Heenan, J.R. Gruen, for the Genes, Reading, and Dyslexia (GRaD) Study Consortium

## Abstract

Previous work has shown that experience speaking more than one language in childhood is associated with decreased intra-individual neural variability in electrophysiological responses during a low-level speech perception task. However, no study has yet evaluated the impact of dual language experience on variability in fMRI responses during a higher-level spoken and written language processing task. In the current study, we calculated trial-by-trial variability in neural activation during an fMRI task that involved deciding whether spoken or printed English words matched pictures of items. We compared trial-by-trial neural activation variability between two groups of 8–15 year-old children: a group of dual language learners (*N* = 24; 11 female) who were Spanish-dominant and acquiring English, and a group of monolingual learners who were English-dominant (*N* = 17; 9 female). We found that when controlling for a variety of language, general cognitive, and demographic measures, neural activation variability for printed words was greater in the dual language learners compared to the monolingual learners in the right middle frontal gyrus, a brain region previously associated with attentional control. This finding highlights how neural variability offers a window of opportunity to examine experience-dependent mechanisms during human development, and motivates future research on bilingual language processing.

## 1. Introduction

Neural variability, an intra-individual measure of fluctuations in brain activity, has been described as a next frontier of human brain mapping that will generate unique insights into cognition and behavior that go beyond those afforded by analyzing average levels of brain activity (Garrett et al., 2013). This claim was initially based on numerous studies showing that neural variability in various brain regions changes in meaningful ways across the lifespan (McIntosh, Kovacevic, & Itier, 2008; Miskovic, Owens, Kuntzelman, & Gibb, 2016; Misic, Mills, Taylor, & McIntosh, 2010; Nomi, Bolt, Ezie, Uddin, & Heller, 2017; Garrett, Kovacevic, McIntosh, & Grady, 2011; Baum & Beauchamp, 2014). More recent work, however, suggests that neural variability may be reflective of additional experiential factors beyond chronological age. In a study examining subcortical and cortical neural variability in response to speech sounds, adolescents who had experience speaking more than one language showed a reduction in the variability of auditory evoked responses compared to their monolingual peers (Krizman, Skoe, Marian, & Kraus, 2014). This reduction in neural variability was correlated with bilingual adolescents’ self report and parental report of proficiency in both English and Spanish. The findings were interpreted to indicate a more robust encoding of speech sounds that was considered a positive consequence of dual language experience (Krizman, Marian, Shook, Skoe, & Kraus, 2012; Krizman et al., 2014). Using a similar experimental paradigm, Skoe and colleagues (2013) also showed that increased maternal education – as a proxy of socioeconomic status (SES) – is similarly associated with a reduction in neural variability.

Despite the observed associations between neural variability and various behavioral and cognitive phenotypes, the parameters that influence the directionality of these associations are still far from clear (Dinstein, Heeger, & Behrmann, 2015). Based on an examination of cortical and subcortical neural variability during different cognitive tasks, previous studies have supported the hypothesis that neural variability within different brain regions can show either increases or decreases, contingent upon multiple factors including the hierarchy in which a brain region is situated as well as the extent to which task performance hinges on cognitive stability versus flexibility (Armbruster-Genc, Ueltzhoffer, & Fiebach, 2016; Misic et al., 2010). In addition to differences in task demands, the nature of the relationship between neural variability and behavior may also differ across imaging modalities based on the temporal and physiological properties of the neural signals that are captured (Centanni et al., 2018). As an example, differences in task and imaging modality may explain why greater reading skills in monolingual children have been associated with reduced variability across trials in brainstem electrophysiological responses to low-level speech sounds (Hornickel & Kraus, 2013), yet the reverse relationship (i.e., increased variability across trials for more skilled readers) has been observed with the fMRI blood oxygen level-dependent (BOLD) signal in cortical regions during a cognitively demanding reading task (Malins et al., 2018).

In light of these mixed findings with respect to task demands, individual differences, and imaging modalities, the current study investigated whether dual language experience modulates fMRI neural activation variability during a reading and language processing task. We adopted the same picture-word matching task used in the Malins et al. (2018) reading study in monolinguals. The reading task, which requires children to decide whether or not printed or spoken words match pictures of items, was chosen for its relatively high cognitive demands. Because of the unique demands of the task, which involves lexico-semantic processing and indexes meta-linguistic knowledge, we hypothesized that dual language experience would be associated with increased neural variability in cortical regions involved in the complex cognitive processes supporting reading, spoken language processing, and attentional control. To test this hypothesis, we compared neural variability between two groups of 8–15 year-old children who differed in their language acquisition history, and had different dominant languages and relative functional usage of English and Spanish. Analyses focused on first quantifying intra-individual trial-by-trial variability in neural activation, and then subsequently evaluating group differences in this intra-individual variability while controlling for potentially confounding factors including age, sex, performance IQ, head motion during scans, and English word reading and vocabulary skills. In addition, regression models included maternal education and family participation in social assistance programs as proxies of SES.

## 2. Methods

### 2.1. Participants

This study was part of a larger, multi-site US and Canadian collaborative project called the Genes, Reading, and Dyslexia (GRaD) Study, with a primary site at Yale University in the US. This project focused on reading development in African American and Hispanic American children to address the absence of large scale studies on the genetics of reading in under-represented populations. The study had a target age range of 8–15 years and followed a case:control design for reading disability. As such, the sample of children in the current study included typically developing children as well as children with reading impairments; however, reading skills were treated continuously in line with recent views on the multifactorial nature of reading disability (Pennington et al., 2012). Exclusion criteria were age outside the target age range; non-minority ethnic or racial group membership; having completed less than three years of English language instruction; foster care placement; preterm birth (< 36 weeks gestation); prolonged stay in neonatal intensive care after birth (> 5 days); history of diagnosed or suspected significant cognitive delays, significant behavioral problems, or frequent school absences; history of serious emotional/psychiatric disturbances (i.e., major depression, psychotic or pervasive developmental disorder, autism spectrum disorder) or a chronic neurologic condition (i.e., seizure disorder, developmental neurological conditions, Tourette’s or other tic disorders, acquired brain injuries); and a documented hearing impairment or vision impairment uncorrected by glasses. Parental consent forms and child assent were collected before participation. This study was approved by the Human Investigation Committee of Yale University and all of the review boards of participating data collection sites.

Children completed a cognitive testing battery that included a number of assessments of reading, language, and intelligence, such as the Woodcock-Johnson III Tests of Achievement (Woodcock, McGrew, & Mather, 2001), the Peabody Picture Vocabulary Test (PPVT-4; Dunn & Dunn, 2007), and the Wechsler Intelligence Scale for Children (WISC-IV; Wechsler, 2003). In addition, each child’s caregiver completed the Disruptive Behavior Rating Scale (Barkley & Murphy, 1998), as well as a questionnaire regarding family background, household resources, health during pregnancy, and their child’s health, educational history, developmental history, and language background.

Neuroimaging data was initially collected from a sample of 100 children who participated in the GRaD study at the Yale site. Of these, data from 61 children passed quality control procedures for task fMRI (i.e., no more than 40% of volumes exceeding the thresholds of .3 mm point-to-point Euclidean movement and/or 10% outlier voxels). Euclidean movement was calculated per volume as the square root of the sum of squares of point-to-point change for each the six motion parameters (i.e., three translation and three rotation).

Using the parental questionnaire, we designated children as having different language acquisition histories and speaking different dominant languages, either English or Spanish. Forty-one children (of the 61 who passed fMRI quality control procedures) met criteria for either the dual language (Spanish-dominant but learning English at school) or monolingual English groups. Dual language (Spanish-dominant) learners (*N* = 24) were defined as children whose parents indicated that Spanish was either the sole primary language spoken in the home, or one of the two primary languages spoken in the home, and their child’s preferred language was Spanish. Monolingual (English-dominant) learners (*N* = 17) were defined as children whose parents indicated that English was the only primary language spoken in the home, their child’s preferred language was English, and their child did not have any formal Spanish instruction. The group affiliation adopted in this study is sensitive to the linguistic context in the United States, where English is the formal language of instruction in the majority of schools, and Spanish is the most representative language among those who do not speak English as their first language (McFarland et al., 2018). If either of the following questions were left unchecked in the parental questionnaire, they were counted as “no” responses: (1) “Is Spanish your child’s primary language?” (2) “When your child started formal school (e.g., Kindergarten or 1st grade), did he or she receive Spanish instruction?”. When designating children as belonging to the different language groups, the twenty children who passed fMRI quality control procedures but did not meet category membership for either the dual language learner or monolingual groups consisted of the following: eight children whose parents indicated that the primary languages spoken in the home were English and Spanish yet their child’s primary language was not Spanish; nine children whose parents indicated that Spanish was the primary language spoken in the home yet their child’s primary language was not Spanish; one child whose primary language in the home was Portuguese; two children whose primary home language could not be properly ascertained because of mismatching responses between the parental questionnaire and an initial screening form filled out by study recruiters. Because these children could not be accurately assigned to one of the two language groups, they were not included in the current set of analyses.

Assessment scores and demographic characteristics concerning the two groups of children are presented in Table 1. As shown in the table, the two groups were matched in age, sex, in-scanner head motion, English word reading, English vocabulary, and the likelihood of meeting criteria for ADHD based on the Disruptive Behavior Rating Scale. The group of dual language learners had a slightly higher mean performance IQ, lower maternal education, a higher likelihood of Hispanic American ancestry, and a marginally higher likelihood of family participation in one or more government assistance programs.

**Table 1.** Assessment scores and demographic characteristics for the two speaker groups.

| Measure | Means of Assessment | Dual language learners<br>(*N* = 24) | Monolingual learners<br>(*N* = 17) | Test of difference between groups |
| --- | --- | --- | --- | --- |
| Age in years | Self-report by the child's caregiver | 10.5 (2.1) | 11.2 (2.0) | $t(35) = -1.09, p = .28$ |
| Sex | Self-report by the child's caregiver | 11 female, 13 male | 9 female, 8 male | $\chi^2(1) = .02, p = .90$ |
| In-scanner motion | Average amount of Euclidean motion across all volumes of scanning (mm/TR) | .15 (.08) | .15 (.10) | $t(29) = -.20, p = .84$ |
| Performance IQ | Wechsler Intelligence Scale for Children (WISC-IV) Matrix Reasoning scaled score | 9.7 (3.0) | 7.5 (2.5) | $t(38) = 2.55, p = .02$ |
| English word reading | Woodcock-Johnson III Letter Word Identification standard score | 94.0 (14.2) | 91.6 (8.7) | $t(38) = .66, p = .51$ |
| English vocabulary | Peabody Picture Vocabulary Test (PPVT-4) standard score | 86.5 (14.1) | 89.3 (12.9) | $t(36) = -.66, p = .52$ |
| Maternal education | Self-reported number of years of formal education | 11.3 (3.1) | 13.9 (1.9) | $t(38) = -3.27, p < .01$ |
| Socioeconomic status | Participation in one or more government assistance programs with income eligibility requirements<sup>1</sup> | 22 indicated participation; 2 did not | 11 indicated participation; 6 did not | $\chi^2(1) = 3.05, p = .08$ |
| Ancestry | Self-report by the child's caregiver | 23 Hispanic American and 1 dual ancestry | 5 Hispanic American, 11 African American, and 1 dual ancestry | $\chi^2(2) = 22.0, p < .001$ |
| ADHD classification | Disruptive Behavior Rating Scale – parent ratings<sup>2</sup> | 4 met criteria for ADHD; 20 did not | 2 met criteria for ADHD; 15 did not | $\chi^2(1) < .01, p = 1$ |

*Note.* Means are listed for all numeric quantities, followed by standard deviations in parentheses.
<sup>1</sup>SES was coded as a binary variable, and was determined by participation in one or more government assistance programs that have income eligibility requirements (Women, Infants, and Children [WIC]; Medicaid/Husky; food stamps; and Section 8 Housing Choice Voucher Program).
<sup>2</sup>Based on DBRS parent ratings, items were first coded using a binary scheme, where 0 (never) or 1 (sometimes) were coded as a 0, and 2 (often) or 3 (very often) were coded as a 1. Missing items were coded as 0. Then, we calculated a sum across the nine items related to inattention, as well as a sum across the nine items related to hyperactivity/impulsivity. Participants were designated as meeting criteria for ADHD if they had a score of six or greater for either inattention, hyperactivity/impulsivity, or both (Lahey et al., 1994; Willcutt et al., 2012).

### 2.2. fMRI Task Procedure

Participants completed a task in which they judged whether a presented picture matched either spoken or printed target words (Frost et al., 2009; Jasińska et al., 2016; Landi et al., 2013; Preston et al., 2015). In this task, participants were presented with pictures of common items (e.g., “plate”) that remained on the screen for 40–65 seconds (7–8 trials) before being replaced by another picture. While each picture remained on the screen, participants were presented with target items in an event-related fashion; specifically, printed words appeared in a box below the picture (presented for 2000 ms in 18-point Verdana font), or spoken words were presented via headphones. The inclusion of printed and spoken words allowed us to examine whether any potential results were specific to print or speech, or were instead more generally related to language processing irrespective of input modality. In one fifth of the trials, the word-picture pairs were matched, while in the other four fifths of trials the word-picture pairs were mismatched. Participants were asked to indicate via button press with their right hand whether or not the printed or spoken word matched the picture. In total, participants completed between four and ten runs of the task with 30 trials per run, six of which were match trials (printed and spoken), and 24 of which were mismatch trials, distributed evenly across the six mismatch conditions (semantically unrelated printed words, semantically related printed words, printed pseudowords, consonant strings, semantically unrelated spoken words, and spoken pseudowords). Therefore, the total number of mismatch trials of each type was between 16 and 40. We only examined responses to printed words and spoken words that were semantically unrelated to picture names to maximize compariability with the Malins et al. (2018) study.

### 2.3. Acquisition of MRI Data

Images were collected using a single 3T TIM-Trio scanner located at the Yale Magnetic Resonance Research Center in New Haven, CT. Anatomical images were acquired in a sagittal orientation (MPRAGE; matrix size = 256 × 256; voxel size = 1 × 1 × 1 mm; FoV = 256 mm; TR = 2530 ms; TE = 2.77 ms; flip angle = 7°). For the picture-word matching task, T2*-weighted images were collected in an axial oblique orientation (25 slices; 6 mm slice thickness; no gap) using single-shot echo-planar imaging (matrix size = 64 × 64; voxel size = 3.44 × 3.44 × 6 mm; FoV = 220 mm; TR = 1550 ms; TE = 30 ms; flip angle = 80°). Participants completed between four and ten functional imaging runs each lasting 3 minutes and 46 seconds (113 volumes). Across all trials in the experiment, the time between trial onsets was jittered between 4 and 13 seconds; trial order and inter-trial intervals (ITIs) were optimized by a MATLAB program developed at Haskins Laboratories that balanced ITIs and null trials across conditions, and minimized the variability of the measured response in Monte Carlo simulations.

### 2.4. Analysis of MRI Data

#### 2.4.1. Preprocessing

Functional images were preprocessed in AFNI (Cox, 1996; RRID:SCR_005927). The first four volumes were removed to allow for stabilization of the magnetic field. Following the removal of these volumes, slice-scan time correction was performed using *3dTshift*. Next, functional images were co-registered with anatomic images, warped to the Colin N27 template in Talairach space using a non-linear transformation (*3dQWarp*), and were corrected for motion by alignment to the beginning of the first functional run (*3dvolreg*). These three steps were concatenated into a single transformation that forced a 3 mm isotropic voxel size on the data. The Colin N27 brain was used for normalization to allow for broader comparison with extant studies, including those with adult samples, and this choice was supported by the finding that relative to the resolution of fMRI data, children over the age of seven show minimal anatomical differences from adults (Burgund et al., 2002). After co-registration, normalization, and motion correction, data were subsequently smoothed using an 8 mm FWHM Gaussian kernel (*3dmerge*), and scaled (*3dcalc*) so that each voxel’s time series had a mean of 100 for each run, with intensity values thereby reflecting percent signal change from the run mean. During this scaling step, values in excess of 200, the default value for scaling in AFNI, were clipped. The default value was selected in order to retain the precision of scaled short values.

#### 2.4.2. GLM Analysis

Next, two different general linear models (GLM) analyses were performed for each participant (*3dDeconvolve*). In the first model, regressors were included for each of the nine stimulus types (pictures, semantically unrelated printed words, semantically related printed words, printed pseudowords, consonant strings, semantically unrelated spoken words, spoken pseudowords, matching printed words, matching spoken words), whereas in the second set of GLMs, single trial regressors were used for each respective condition of interest (i.e., either semantically unrelated printed words or semantically unrelated spoken words) while all other conditions were kept as condition-wise regressors (using the flag *-stim_times_IM* in *3dDeconvolve* in AFNI for the condition with single trial regressors). Then, we calculated the difference in the variance of the residuals between the GLM *with* trial-wise regressors and the GLM *without* trial-wise regressors to compute a difference-of-residuals map (Armbruster-Genc et al., 2016). This procedure was respectively performed for the semantically unrelated printed mismatch and semantically unrelated spoken mismatch conditions. In all GLMs, a two-parameter SPM gamma variate basis function (‘SPMG2’ in *3dDeconvolve*) was used to approximate the hemodynamic response function (HRF), with temporal derivatives included to allow for deviations from the canonical HRF (Henson, Price, Rugg, Turner, & Friston, 2002). In addition, all GLMs included regressors for the six motion parameters and second-order polynomial drift terms for each run, and volumes were censored if they exceeded the threshold of .3 mm point-to-point Euclidean movement and/or 10% outliers.

#### 2.4.3. Groupwise Statistical Analysis

Group-wise analysis was performed in AFNI using a multivariate model (*3dMVM*) with difference-of-residual maps as the dependent variable (for either printed or spoken words), and the following between-subjects variables: language group (dual language learners versus monolingual learners), age in months, sex, average amount of Euclidean motion per volume of scanning, performance IQ (WISC Matrix Reasoning raw scores), English word reading scores (raw scores from the Letter Word Identification subtest from the Woodcock-Johnson III Tests of Achievement), English vocabulary (raw scores from the Peabody Picture Vocabulary Test), number of years of maternal education, and socioeconomic status (SES). Note that because we included age as a covariate, we elected to use raw rather than standard scores for each of the three assessment measures. SES was coded as a binary variable, and was determined by participation in one or more government assistance programs that have income eligibility requirements (Women, Infants, and Children [WIC]; Medicaid/Husky; food stamps; and Housing Choice Voucher Program, Section 8). For these group-wise analyses, the voxel-wise threshold was *p* = .005, cluster corrected at *p* = .05. Cluster correction was performed by estimating the spatial autocorrelation of the residual time courses for each subject (*3dFWHMx*), and then using mean parameter values across subjects as inputs to the simulation program *3dClustSim* (10,000 iterations). This procedure was based on the latest recommendations for cluster correction in fMRI (Eklund, Nichols, & Knutsson, 2016). The cluster threshold for an alpha level of .05 was 143 voxels.

## 3. Results

As shown in Figure 1, we observed greater neural activation variability for printed word trials in the group of dual language learners compared to the monolingual learners in a cluster located in the right middle frontal gyrus (MFG), extending into the inferior frontal gyrus (peak voxel 23, 2, 39 in Talairach space; cluster extent of 205 voxels with a corrected p-value of .02). As shown in the boxplot in Figure 1, one observation in the dual language learner group was an outlier, which was confirmed by the program *outlierTest* in version 3.0–2 of the ‘car’ package (Fox & Weisberg, 2011) of the R Project for Statistical Computing (R Core Team, 2017; RRID:SCR_00195). This outlier can be considered an influential observation (Bonferonni *p* = .002); therefore, Table 2 displays the results of the multiple regression analysis that was performed using the program *lm* in version 3.4.1 of the ‘stats’ package in R, after this outlier was removed. As displayed in the table, other variables beyond dual language experience influenced neural activation variability: namely, amount of head motion, sex, and SES. The direction of each of these relationships was as follows: neural activation variability increased as a function of head motion, was reduced in males compared to females, and was reduced in low relative to high SES environments. It should be noted that even with the outlier present in the model, the effect of language group remains significant (*p* < .001).

**Figure 1.**
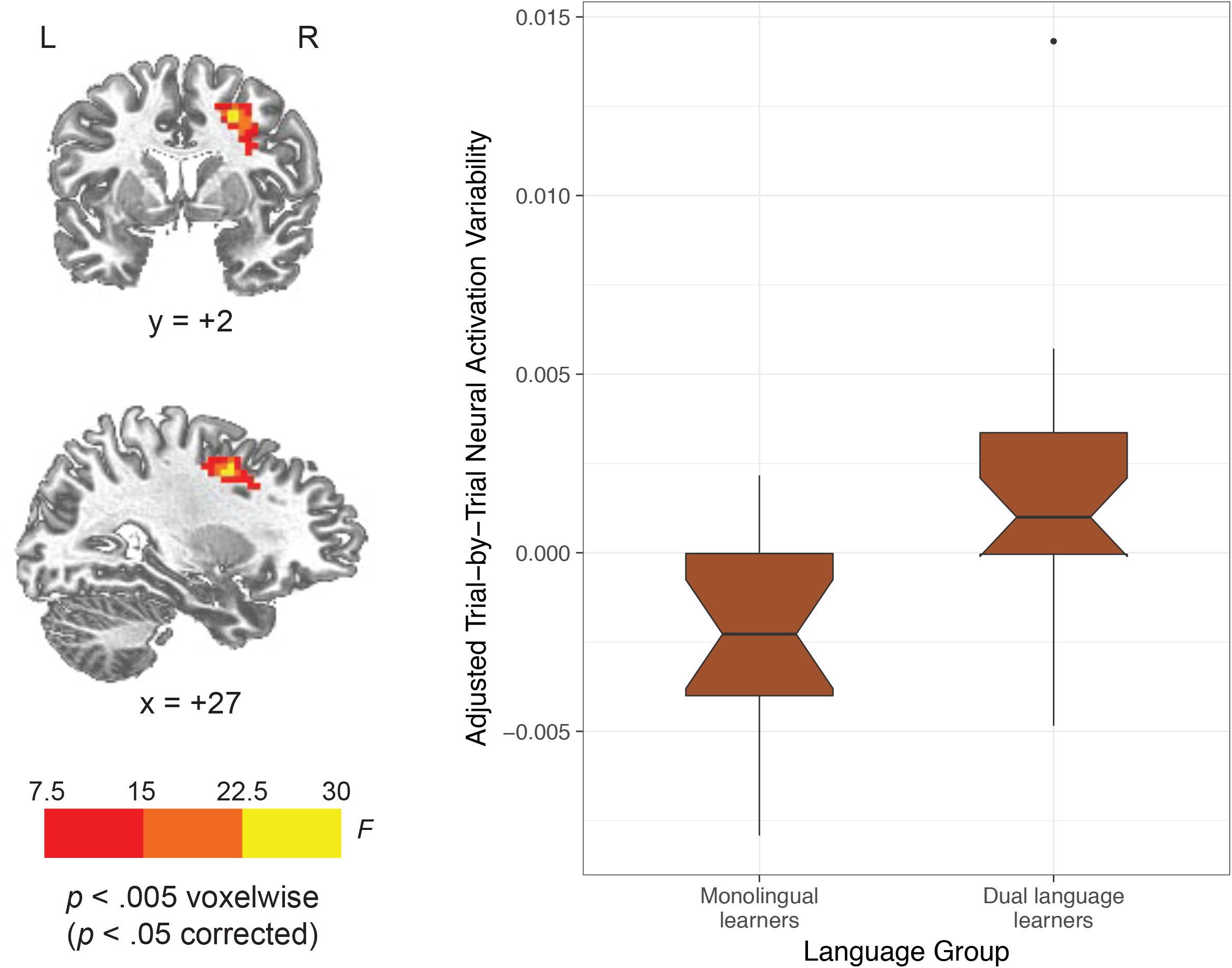
The cluster in the right middle frontal gyrus that showed a main effect of language group on trial-by-trial neural activation variability for printed words, when adjusting for the following covariates: age, sex, head motion, performance IQ, English word reading skills, English vocabulary, maternal education, and SES. The boxplot shows the distribution of adjusted trial-by-trial variability values within this cluster for each of the two language groups.

**Table 2.** Multiple regression predicting signal in the right middle frontal gyrus from the analysis of all printed word trials in the full sample, with one outlier participant removed (*N* = 40).

| Regressor | *B* | *SE* | *t* | *p* |
| --- | --- | --- | --- | --- |
| Language category (1=dual language learners) | .00482 | .00112 | 4.292 | <.001 |
| Age in months | .00000 | .00003 | 0.066 | .948 |
| Sex (1=male) | -.00353 | .00086 | -4.088 | <.001 |
| Average Euclidean movement (mm/TR) | .01876 | .00491 | 3.820 | .001 |
| Performance IQ (WISC Matrix Reasoning raw scores) | -.00014 | .00011 | -1.307 | .201 |
| English word reading (Woodcock-Johnson III Letter Word Identification raw scores) | -.00001 | .00007 | -0.143 | .887 |
| English vocabulary (PPVT raw scores) | -.00001 | .00003 | -0.201 | .842 |
| Maternal education in years | .00030 | .00016 | 1.814 | .080 |
| Socioeconomic status (1=low) | -.00247 | .00116 | -2.132 | .041 |

When we examined neural activation variability for spoken word trials, no brain regions showed significant differences in variability between the two groups. In addition, to test the specificity of the effect, we also repeated all analyses for both printed and spoken word trials using measures of mean activation variability instead of neural activation variability. This was done by performing a standard GLM analysis in which all trials within each condition type were modeled using a single regressor as opposed to separate regressors for each individual trial. These analyses did not reveal any significant differences between the two groups in mean activation for either printed or spoken word trials.

One limitation of this analysis is that differences between groups in neural activation variability might be attributable to potential differences in task performance. Unfortunately, due to technical issues, in-scanner behavioral data was missing from 12 of the 41 participants. In addition, for five additional participants, valid responses were recorded for less than sixty percent of all trials. For the remaining 24 participants, we calculated non-parametric estimates of sensitivity and bias (Stanislaw & Todorov, 1999), as was done in Frost et al. (2009), which also employed the picture-word matching task used in the current study. Sensitivity (A′) was high across all participants (mean = .97; SD = .02), whereas bias (B″) was fairly low (mean = .09; SD = .59), indicating that most children understood and could perform the task. When calculating sensitivity and bias, trials with outlier reaction times were ignored. These trials were defined as those with reaction times less than 200 ms or greater than 1.5 times the interquartile range above the third quartile of a participant’s distribution of reaction times. Sensitivity did not differ between the dual language learners and monolingual learners, even when accounting for age, sex, performance IQ, amount of head motion, English vocabulary, English word reading scores, maternal education, and SES (*B* = .008, *SE* = .012, *p* = .51). However, bias was significantly lower in the dual language group compared to the monolingual group when controlling for this same set of covariates (*B* = −1.04, *SE* = .396, *p* = .02).

To rule out differences in task performance, we coded trials as either correct or incorrect and re-performed GLM analyses with correct trials only for the subset of 24 participants from whom we had usable in-scanner behavioral data. In these models, all incorrect trials and all trials with outlier reaction times, regardless of condition type, were collapsed into a single regressor.

Using this approach, we computed difference-of-residuals maps for correct trials only, and then performed the same multiple regression analysis that was implemented with the larger sample for the cluster in the right MFG. As displayed in Table 3, we observed similar relationships in this subset of 24 participants as we did in the larger sample; namely, dual language experience was associated with increased neural activation variability in the right MFG, whereas low SES was associated with decreased neural activation variability in this same region.

**Table 3.** Multiple regression predicting signal in the right middle frontal gyrus from the analysis of correct printed word trials from participants with usable in-scanner behavioral data (*N* = 24).

| Regressor | *B* | *SE* | *t* | *p* |
| --- | --- | --- | --- | --- |
| Language category (1=dual language learners) | .00436 | .00150 | 2.899 | .012 |
| Age in months | -.00005 | .00003 | -1.863 | .084 |
| Sex (1=male) | -.00102 | .00113 | -.904 | .381 |
| Average Euclidean movement (mm/TR) | .01465 | .00918 | 1.596 | .133 |
| Performance IQ (WISC Matrix Reasoning raw scores) | -.00006 | .00013 | -.463 | .651 |
| English word reading (Woodcock-Johnson III Letter Word Identification raw scores) | .00005 | .00008 | .627 | .541 |
| English vocabulary (PPVT raw scores) | .00005 | .00003 | 1.398 | .184 |
| Maternal education in years | .00046 | .00027 | 1.731 | .105 |
| Socioeconomic status (1=low) | -.00380 | .00121 | -3.126 | .007 |

## 4. Discussion

Our aim was to assess whether dual language experience modifies neural activation variability during an fMRI reading and language task that involves deciding whether or not spoken and printed words match pictures of items. This was motivated by recent work documenting differences in bilingual compared to monolingual adolescents in subcortical and cortical neural variability to speech sounds as measured using auditory evoked responses (Krizman et al., 2014), as well as studies exploring the relationship between neural variability and reading skill in monolingual children (Centanni et al., 2018; Hornickel & Kraus, 2013; Malins et al., 2018). We hypothesized that, similar to what has been observed with respect to reading skill, although dual language experience has been associated with a reduction in neural variability during lower-level perceptual tasks, in this higher-level cognitively engaging task, dual language experience would instead be associated with increased variability in the BOLD signal in cortical regions involved in the complex cognitive processes supporting reading, spoken language processing, and attentional control.

### 4.1. The Effect of Dual Language Experience on Neural Activation Variability

For the printed word trials, dual language learners showed greater variability in neural activation compared to their monolingual peers in the right middle frontal gyrus. We consider the task to be meta-linguistic in nature, in that it requires participants to actively engage lexico-semantic processing systems to decide whether the printed or spoken input matches expectations generated by a picture. In addition, this active mode of processing places a considerable degree of attentional demands on participants. Given these high demands on reading, language, and attentional control systems, the task may have promoted cognitive flexibility as opposed to stability, and thereby drove neural variability to increase (Armbruster-Genc et al., 2016). This could explain why the directionality of the effect is similar to what has been observed for reading in monolinguals: even though dual language experience is associated with reduced neural variability in a lower-level task such as passive listening to speech sounds (Krizman et al., 2014), it is associated with increased neural variability during a higher-level meta-cognitive task. However, this finding should be interpreted with the caveat that in addition to task differences, the current measure of trial-to-trial BOLD signal variability differs from variability in auditory evoked responses both temporally (on the order of seconds/minutes as opposed to milliseconds) and physiologically (hemodynamic as opposed to electrophysiological responses).

In this particular task context, the dual language learners faced additional demands compared to monolingual learners because they were processing the stimuli in their non-dominant language, which may have resulted in more extensive engagement of attentional control regions in the brain (Marian, Bartolotti, Rochanavibhata, Bradley, & Hernandez, 2017). In a previous fMRI study in which children were presented with English speech sounds, middle frontal regions showed stronger engagement in nine to ten year-old Spanish-English bilingual children compared to English monolingual children of the same age, and these differences were thought to reflect stronger recruitment of higher-order executive areas when processing non-native speech (Archila-Suerte, Zevin, Ramos, & Hernandez, 2013). The current finding of increased neural activation variability in this region for the dual language learners when reading in their non-dominant language complements this result. However, we only observed the effect for printed words and not for spoken words, suggesting that the unique demands of reading may underlie the observed results. Additionally, the lack of a language difference between groups for spoken word trials might be attributable to imaging modality: for example, Centanni et al. (2018) speculate that perhaps because of the fine temporal structure of speech sounds, a temporally sensitive technique such as magnetoencephalography may be more likely than fMRI to detect differences in neural variability when individuals are processing speech.

A noteworthy finding is that in the current study, differences between language groups were observed in neural activation variability rather than in mean activation. Amidst concerns regarding the locus and consistency of the effects of bilingual language experience on behavior (Paap, Johnson, & Sawi, 2015), the current findings suggest that analyses of neural activation variability may constitute an innovative approach to measuring the effects of bilingual language experience on brain activity, which may help clarify outstanding issues in the field of bilingual research (van Heuven & Coderre, 2015).

One limitation of the current study is that task performance data was missing from a number of the participants due to technical issues. When we analyzed task performance in the participants from whom we had usable data, sensitivity did not differ between the two groups, although bias was lower in the dual language group. When we re-performed analyses for the right MFG using correct trials only, we observed a similar pattern of results as we did for the full sample. For this reason, we assert that differences between groups cannot be entirely attributed to potential differences in task performance.

### 4.2. Socioeconomic Influences on Neural Activation Variability

Besides dual language experience, an important factor that influenced neural activation variability was SES. Although examining neural activation variability in relation to SES was not a primary aim of this study, we included this variable as a covariate because of previous work showing that socioeconomic factors such as parental education are associated with brain development and brain-behavior relationships related to reading (Hackman & Farah, 2009; Noble et al., 2015; Noble, Wolmetz, Ochs, Farah, & McCandliss, 2006), as well as decreased neural response consistency to speech sounds (Skoe, Krizman, & Kraus, 2013), and furthermore can interact with dual language experience to influence brain structure and cognition (Brito & Noble, 2018). The current pattern of results was such that lower SES was associated with a reduction in neural activation variability within the same cortical region that showed enhancements in relation to dual language experience. Because 33 out of the 41 children, or 80% our sample, came from low SES environments, this finding supports the recent claim that enhancements in neural functioning associated with dual language experience are not restricted to high SES environments (Krizman, Skoe, & Kraus, 2016). Although we did not have data from enough dual language learners with high SES to test for an interaction between dual language experience and SES, given the recent report of an interaction between these two variables in terms of brain structure (Brito & Noble, 2018), future functional MRI studies with larger cohorts should aim to elucidate the extent to which language experience and SES interact to drive changes in neural variability in cognitive systems during development.

## 5. Conclusions

The current study suggests that dual language experience modulates neural activation variability during a reading and language processing task that places considerable demands upon reading, language, and attentional control systems. This finding has implications for bilingual language research, as changes in neural variability could be one of the ways by which experience speaking more than one language shapes different cognitive systems, such as those involved in attentional control (Bialystok, Craik, & Luk, 2012; Costa & Sebastián-Gallés, 2014; Krizman et al., 2012; Krizman et al., 2014). Beyond its implications for bilingual language research, these results also suggest that experiential factors such as speaking more than one language, as well as living in a low SES environment, not only influence electrophysiological response variability during a low-level speech perception task (Krizman et al., 2012; Krizman et al., 2014; Skoe et al., 2013), but can also affect BOLD signal variability during a higher-level reading and language processing task.

There is some evidence that biological factors, such as genetic variants of the dyslexia susceptibility gene *KIAA0319* (Centanni et al., 2018), as well as dopamine receptor density in subcortical and cortical regions (Guitart-Masip et al., 2016), influence neural variability in different populations. It is our view that future studies should explore how these biological factors interact with different childhood factors, including dual language experience and SES, to dynamically shape neural variability within a developmental context (e.g., Hernandez, Claussennius-Kalman, Ronderos, & Vaughn, 2018; Hernandez, Greene, Vaughn, Francis, & Grigorenko, 2015; Vaughn et al., 2016; Vaughn & Hernandez, 2018). It will be especially important to understand how neural variability changes in relation to specific experiences in a child’s life, as the malleability of neural variability may have important implications for cognitive performance in children across various academic domains.

## Acknowledgments

Research reported in this publication was supported by The Manton Foundation as well as the Eunice Kennedy Shriver National Institute of Child Health & Human Development of the National Institutes of Health under award number P50HD027802. The content is solely the responsibility of the authors and does not necessarily represent the official views of the National Institutes of Health. The authors do not have any conflicts of interest to declare.

